# An Approach to Measuring Protein Turnover in Human Induced Pluripotent Stem Cell Organoids by Mass Spectrometry

**DOI:** 10.1101/2021.09.30.462679

**Authors:** Anthony Duchesne, Jing Dong, Andrew N. Bayne, Nguyen-Vi Mohamed, Wei Yi, Meghna Mathur, Edward A. Fon, Thomas M. Durcan, Jean-François Trempe

## Abstract

Patient-derived organoids from induced pluripotent stem cells have emerged as a model for studying human diseases beyond conventional two-dimensional (2D) cell culture. Briefly, these three-dimensional organoids are highly complex, capable of self-organizing, recapitulate cellular architecture, and have the potential to model diseases in complex organs, such as the brain. For example, the hallmark of Parkinson’s disease - proteostatic dysfunction leading to the selective death of neurons in the substantia nigra - present a subtle distinction in cell type specificity that is simply lost in 2D cell culture models. As such, the development of robust methods to study global proteostasis and protein turnover in organoids will remain a critical need as organoid models evolve. To solve this problem, we have designed a workflow to extract proteins from organoids and measure global protein turnover using mass spectrometry and stable isotope labeling using amino acids in cell culture (SILAC). This allowed us to measure the turnover rates of 844 proteins and compare protein turnover to previously reported data in primary cell cultures and *in vivo* models. Taken together, this method will facilitate the study of proteostasis in organoid models of human disease and will provide an analytical and statistical framework to measure protein turnover in organoids of all cell types.

## 1.1 Organoids as a novel model to recapitulate neurodegenerative disease

Three dimensional (3D) human brain organoids derived from induced pluripotent stem cells (iPSCs) have emerged as a novel tool in modelling distinct regions of the brain and can even reconstitute neuronal crosstalk via organoid fusions [1–4]. This technology serves as a critical bridge between 2D cultures and *in vivo* models in examining complex neural mechanisms and their dysregulation in disease. Unlike traditional *in vitro* cultures, the architecture of organoids consists of multiple region-specific cell types conferring more physiologically relevant characteristics [5,6]. Two examples of disease-relevant hallmarks in organoids that remain poorly captured in 2D culture models are as follows: (1) midbrain organoids are capable of producing neuromelanin-like granules, a distinct structure resulting from dopamine synthesis that are highly enriched in the neurons lost in Parkinson’s disease (PD) [1,5,7], and (2) β-amyloid plaques and neurofibrillary tangles are found in cerebral organoids, a pathological marker of Alzheimer’s disease (AD) pathology [8].

These findings demonstrate that brain organoids complement existing model systems as a tool to study the mechanisms underlying neurodegenerative diseases. However, despite their promise as a driver for scientific discovery, the technology is still in its infancy as validated approaches focusing on experimental methods to study these models are lacking.

## 1.2 Protein Dynamics in Neurodegeneration

Turnover, the dynamic process of the removal and replacement of proteins, is essential to maintaining the homeostasis of all cells including neurons [9]. Studies have shown that dysfunctional mitochondria and their impaired turnover is a fundamental problem associated with specific neurodegenerative disorders such as PD, AD and Amyotrophic Lateral Sclerosis [10]. In fact, mutations in PINK1 and Parkin, two proteins implicated in the selective turnover of mitochondria, cause autosomal recessive juvenile PD [11]. Furthermore, defects in both the ubiquitin proteasome system and autophagy lead to protein misfolding and aggregation, a common mechanism of pathogenesis in neurodegenerative diseases, such as PD, AD and Huntington’s Disease [12,13]. As such, the proteome-wide study of protein dynamics in organoids presents a unique opportunity to uncovering novel mechanisms of neurodegeneration. Other studies have used quantitative mass spectrometry to profile differential protein expression in brain organoids following drug treatment [14,15]. However, there are currently no established methods to measure protein turnover in these systems. Our study aims to address that gap and provides a robust methodology for protein turnover measurement in organoids.

## 1.3 Measuring protein turnover in organoids using SILAC

Protein turnover can be measured using stable isotope labelled amino acids (SILAC) coupled with quantitative proteomics by mass spectrometry (MS). While stable isotope labels can be introduced into proteins either metabolically, chemically, or enzymatically, this study will focus on metabolic labelling, as it is the most effective implementation for *in-* and *ex-vivo* systems [16]. Briefly, the metabolic labelling approach of SILAC involves growing cells in two separate media: (1) the “light” medium, which contains an amino acid with all atoms at their natural isotope abundance and (2) a “heavy” medium, which contains heavy isotope labels incorporated into the same amino acid. The “heavy” labeled amino acid is subsequently incorporated into newly synthesized proteins, which induces a small mass shift in the digested peptides that is distinguishable by MS. The ratio of heavy to light (H:L) abundance of each peptide can be measured at different time-points in a pulse-chase-like time course experiment. The H:L ratios for all peptides of a given protein can then be averaged to compute a half-life for that particular protein. One critical requirement for the measurement of turnover in this manner is that the protein levels must remain constant throughout the time course (steady state); in this case, the rate of synthesis must equal the rate of degradation, allowing the turnover rate to be calculated. Common SILAC labels apply ^13^C or ^15^N isotopes within Arg or Lys in media to 2D cell culture systems, or in heavy labelled food of animals such as zebrafish, newts and mouse. [17–21]. ^13^C_6_-labeled Lys, an essential amino acid, has been used to measure the turnover of proteins in mice [22]. Alternatively, leucine is also an essential amino acid that is highly abundant, and less costly than its Lys/Arg label counterparts. Furthermore, leucine does not undergo metabolic scrambling, the process in which the heavy label is metabolized and incorporated into other amino acids potentially confounding analysis [23]. Leucine is indeed catabolized to a-ketoisocaproic acid and β-hydroxy-β-methylbutyric acid, two metabolites that enter cholesterol biosynthesis or the citric acid cycle via acetyl-CoA, where the branched aliphatic δ carbons are excreted through carbon dioxide.

Other studies have validated the use of heavy leucine in turnover measurements by feeding *Drosophila melanogaster* [5,5,5]-deuterium-3-leucine (D3-Leu) food that was incorporated into the flies over time [24]. Likewise, organoid models can be cultured with ^13^C,^15^N-labeled lysine and arginine to characterize growth and protein abundance under different conditions [25]. Here, we report a robust proteomic method that measures the half-life of proteins in iPSC-derived organoid tissue using D3-Leu as a tracer. Our approach has been optimized to produce robust and consistent data from a variety of protein processing and mass spectrometry methods.

## 2. Methods

### Cell-line information and ethical approvals

The use of iPSCs in this research is approved by the McGill University Health Centre Research Ethics Board (DURCAN_IPSC / 2019-5374). AIW002 lines come from C-BIG repository, The Neuro.

### 2.1 Organoid Generation

Human midbrain organoids (hMOs) derived from healthy individuals were provided by the Neuro’s Early Drug Discovery Unit (https://www.mcgill.ca/neuro/open-science/eddu). The complete procedure regarding generation is described in the standardized protocol ref: https://doi.org/10.12688/mniopenres.12816.2 [26].

### 2.2 In Vivo Stable Isotope Labeling of hMOs

The SILAC time course experiment consisted of triplicate hMOs (n = 3) at five different time points: Day 0, 3, 7, 14 and 28. Sixty-day old hMOs were incubated in light SILAC media (Tables 1 and 2). After 7 days of incubation, 3 hMOs were extracted as day 0 (baseline), flash frozen and stored at −80°C until needed. The remaining hMOs were transferred to heavy SILAC media and left to incubate for the corresponding number of days. Both heavy and light media were changed every 2-3 days. All other hMOs were then extracted, frozen, and stored at each designated timepoint following the time course schedule. As a negative control for heavy isotope incorporation, 3 hMOs were also grown in light media and were harvested on Day 28. These hMOs were not included in any turnover calculations.

**Table 1:**
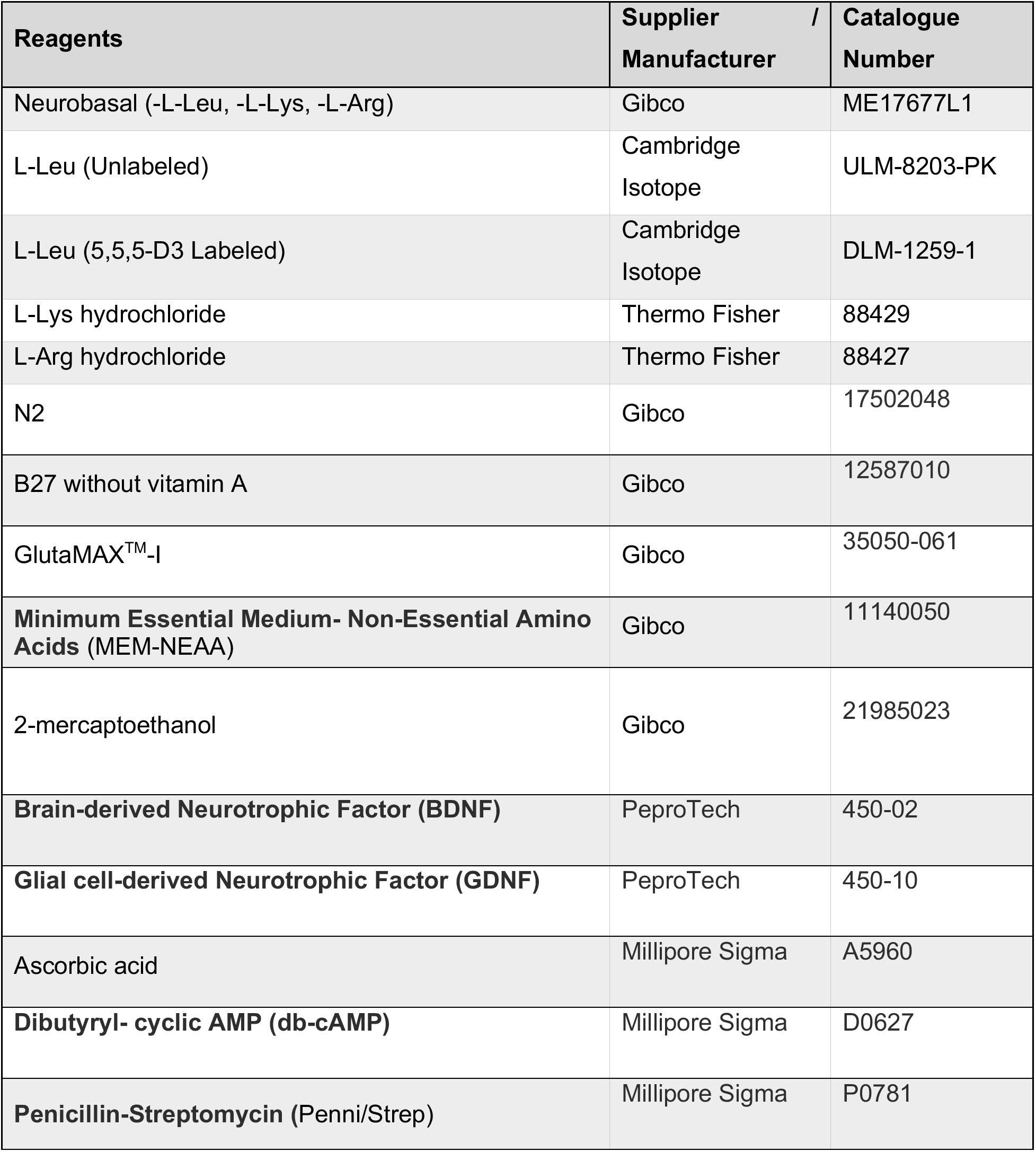
Manufacturer information regarding media and biochemical reagents used for organoid labeling

**Table 2:** Recipe for preparation of labeling medium

| Reagent and Final Concentration | Recipe for 50 mL |
| --- | --- |
| Neurobasal (without L-Leu, L-Lys or L-Arg) | 50 mL |
| 1:100 N2 | 0.5 mL |
| 1:50 B27 without vitamin A | 1 mL |
| 1% GlutaMAX™-I | 0.5 mL |
| 1% MEM-NEAA | 0.5 mL |
| 2-mercaptoethanol | 0.175 µL |
| 10 ng/mL BDNF | 25 µL |
| 10 ng/mL GDNF | 25 µL |
| 100 µM ascorbic acid | 25 µL |
| 125 µM db-cAMP | 12.5 µL |
| Penni/Strep | 0.05 mL |
| 105 mg/L *L-Leu (Unlabeled) / L-Leu (5,5,5-D3 Labeled) | 0.5 mL |
| 146 mg/L L-Lys | 0.5 mL |
| 84 mg/L L-Arg | 0.5 mL |

### 2.3 hMOs Sample Preparation

hMOs were removed from −80°C storage and rinsed in buffer (50 mM Tris • HCl, pH 7.5). They were then placed in a Potter-Elvehjem PTFE glass tube Dounce homogenizer with 200 μL of lysis buffer (50 mM Tris • HCl, pH 7.5, 8 M urea, 1 mM EDTA, 1X Halt™ Protease Inhibitor Cocktail, 1X PhosStop™ Phosphatase Inhibitor Cocktail). Each hMO was homogenized in the lysis buffer with 30-35 pestle strokes and transferred to a 1.5 mL Eppendorf. The tubes were sonicated in a bath sonicator for 10 minutes and spun at 16,000 g for 10 minutes at 4°C. The supernatants were collected and placed into new low-bind 1.5 mL tubes on ice. Protein concentrations of each supernatant were measured using Pierce BCA protein assay kit, according to manufacturer’s instructions. All samples were normalized to 1 μg /μL in 1X Laemmli buffer and were boiled at 80°C for 5 minutes for subsequent SDS-PAGE in-gel digestion.

### 2.4 In-Gel Digestion

For each sample, 20 μg of organoid lysates were loaded onto a 10% Mini-PROTEAN® TGX™ Precast Protein Gels (50 μl wells). Samples were run at 100 V until the samples migrated fully into the stacking region of the gel. Protein bands were visualized using SimplyBlue™ SafeStain (ThermoFisher), according to manufacturer instructions. Each sample was excised in a single band using a clean razor blade, and in-gel digestion was performed as previously described [27]. Briefly, each band was destained, reduced with 10 mM DTT, alkylated with 55 mM iodoacetic acid, and digested with trypsin overnight at 37 °C. Digested peptides were extracted with 1:2 (vol/vol) 5% formic acid / acetonitrile, transferred to a clean 1.5 mL Eppendorf tube, and dried in a Savant SPD2010 SpeedVac (ThermoFisher). Peptides were re-suspended in 0.1 % formic acid, and their concentrations were measured using the Pierce Quantitative Colorimetric Peptide assay.

### 2.5 LC-MS/MS

2 ug of extracted peptides were re-solubilized in 0.1% aqueous formic acid / 2% acetonitrile and loaded onto a Thermo Acclaim Pepmap (Thermo, 75 μm ID X 2 cm C18 3 μm beads) pre-column and then onto an Acclaim Pepmap EASY-Spray (Thermo, 75 μm X 15 cm with 2 μm C18 beads) analytical column separation using a Dionex Ultimate 3000 uHPLC at 250 nl/min with a gradient of 2-35% organic (0.1% formic acid in acetonitrile) over 3 hours. Peptides were analyzed using a Thermo Orbitrap Fusion mass spectrometer operating at 120,000 resolution (FWHM in MS1) with HCD sequencing (15,000 resolution) at top speed for all peptides with a charge of 2+ or greater.

### 2.6 Data Processing

Our data processing protocol is easily portable to data generated from different mass spectrometer vendors and/or digestion methods, as it utilizes freely available, open-source software. The basic workflow consists of: (1) peptide identification from raw data files in MaxQuant; (2) spectral library building in Skyline; (3) protein half-life determination in Topograph; (4) data parsing, filtering, and analysis.

#### 2.6.1 Peptide identification and database search with MaxQuant

RAW mass spectra data was processed using Andromeda, integrated into MaxQuant (version 1.6.5) [28]. While MaxQuant has the ability to specify heavy labels and calculate H:L peptide ratios directly, our workflow uses MaxQuant solely as a means for protein identification for spectral library building in Skyline.

1. Load all RAW data into MaxQuant using “Load folder” with “Recursive” selected, and give each file a name with “Set experiment”. Biological replicates must have unique experiment names (eg. run_1, run_2, run_3), as they will be combined later in Topograph.
2. In “Group-specific parameters”, select carbamidomethylation (C) as a fixed modification. Select oxidation (M) and protein acetylation (N-term) as variable modifications. For instrument parameters, select default MaxQuant parameters for an Orbitrap, including a first search peptide tolerance of 20 ppm and a main search peptide tolerance of 4.5 ppm.
3. Select Trypsin/P as an enzyme for cleavage, and permit a maximum of two missed cleavages.
4. In “Global parameters”, add your FASTA file of interest (ie. reviewed human proteome from UniProt; UP000005640). Select the row and set the identifier rule to “Uniprot identifier”. The minimum peptide length can be left at 7 a.a.
5. In “Identification”, ensure that you select “Match between runs” to enable transferring of protein identifications across runs. All other settings can be left default.
6. Set the number of dedicated processors in the bottom left and start the run.

#### 2.6.2 Building Spectral Library through Skyline

Skyline (version 21.1) is an open-source application for targeted proteomics and quantitative data analysis [29]. Skyline can build spectral libraries, collections of known peptide sequence spectra, which are then used to identify and compare unknown mass spectra. For additional details and tutorials, visit the Skyline website: https://skyline.ms/project/home/software/Skyline/begin.view

1. Save RAW files and all MaxQuant output files in the same directory.
2. Create a blank Skyline document and save it the same folder with the RAW data and MaxQuant text file outputs
3. Select ‘File’ to import a ‘Peptide Search’. Navigate to the msms.txt file and import with default settings.
4. Select the same FASTA file used for the MaxQuant search and import RAW files to create a BiblioSpec spectral library.

#### 2.6.3 Protein Half-Life Calculations with Topograph

Topograph is able to process spectral libraries to calculate protein turnover rates through analyzing the fraction of heavy labels in newly synthesized proteins. The software is able integrate information from all biological replicates across all the timepoints to produce a half-life of a given protein. Furthermore, Topograph takes into account precursor pool enrichment levels, allowing for accurate calculations when the precursor pool is not fully labeled [30].

1. Create a new workspace in the same directory as the BiblioSpec library and RAW data files.
2. Navigate to ‘Add Search Results’ to select ‘Import BiblioSpec library’
3. Keep the default static modification (C heavier by 57.021461 Da) and specify the heavy label by selecting the preset ‘D3-Leu’ option. Custom isotope labels can also be configured, if necessary.
4. Select and import RAW data files and begin analysis on peptides with default settings. This process can take several days depending on the complexity, quantity and size of the data.
5. After the peptide analysis is complete, select ‘Set Cohort and Time points of Samples’ and assign time points and the cohort of samples based on the experimental design. Specify the number of biological replicates and conditions associated with the experiment.
6. Prior to calculating half-lives, configure the following parameters under ‘View half-lives’: Select the option for ‘Distribution of Unlabeled, Partially Labeled and Fully Labeled Peptides’. Set the percent of label at the start of the experiment to 0 and choose the median precursor pool. Set a minimum intensity of 10^5^, minimum deconvolution score of 0.95, minimum turnover score of 0.98 and an outlier filter of TwoStdDev for the acceptance criteria. Choose ‘Simple Linear Regression with 95% CI’ for further statistical analysis. The curve should not be forced through the origin as there is a time delay from the introduction of the label to the appearance of the label in the peptide.
7. To calculate half-lives, select ‘By Protein’, then click ‘Recalculate’
8. Select the view tab and navigate towards the options “Half Lives” and “Results By Replicate” to output “ResultRow” and “PerResultReplicate” as csv files. “ResultRow” is a table listing all the identified proteins and their corresponding half-lives and confidence intervals. “PerResultReplicate” is an overview of all the individual peptides found in each RAW file and displays the heavy label incorporation on a peptide level.

#### 2.6.4 Data parsing, Filtering and Analysis

Data cleansing is conducted through a combination of an in house-implementation of Excel VBA macros and manual validation.

1. Using the ‘PerResultReplicate.csv’ file, remove proteins with less than 2 peptides and less than 15 data points that contribute to the half-life calculation.
2. Using the ‘ResultRow.csv’ file, remove all proteins that have ‘NA’ values for their half-life or 95%confidence interval.
3. Divide the range of the 95% confidence interval with the half-life of each individual protein to yield a value analogous to the coefficient of variation. Exclude proteins with a “coefficient of variation” ratio of >0.3.

### 2.7 Protein Abundance Calculations

Topograph sums the peak areas of all forms of both the labelled (heavy) and unlabelled (light) peptide as a measure of total abundance. This abundance should be compared across time-points, either on a peptide or a protein level, to ensure that the steady state assumption remains valid.

1. Normalize the abundance values under the ‘Area’ column found in the ‘PerResultReplicate.csv’ by dividing each individual value by the sum of all the values in that biological replicate.
2. Calculate the average abundance values for each peptide across all the biological replicates at day 0 and 28.
3. For each peptide, perform paired sample t-tests with a Benjamini Hochberg correction to compare the mean abundance values between day 0 and 28.
4. Exclude proteins associated with peptides that have significant differences in mean abundance from further turnover analysis.

### 2.8 Statistical Analysis and Annotation

All analyses and the generation of figures were performed through GraphPad Prism (GraphPad Software). Proteins were assigned a functional annotation using information from a variety of resources including gene and protein information databases (MitoCarta, COMPARTMENTS and KEGG) [31–33]. Functional enrichment analysis was performed with STRING (version 11.5), a database to predict and visualize protein interaction networks [34].

## 3.1 Characterizing Protein Half-Lives

D3-Leu was successfully incorporated into hMOs following incubation with D3-Leu media, which can be visualized in the MS1 mass spectrum by a corresponding 3 Da rightward shift for a given peptide (*ie*. 1.5 m/z shift for a peptide of 2+ charge) (Figure 2A). The absence of heavy label peaks in both D0 heavy and D28 light media only conditions confirm the selective incorporation of D3-Leu and highlight the robustness of D3-Leu based quantification. The hMOs also continued to incorporate heavy labels over time, as shown by increasing H:L ratios for peptides at later time points (Figure 2B). Specific protein turnover curves and half-life calculations were generated in Topograph. In the example provided, the electron transport chain protein ATP5A yielded a half-life of 14 ± 0.28 days (Figure 2C).

**Figure 1:**
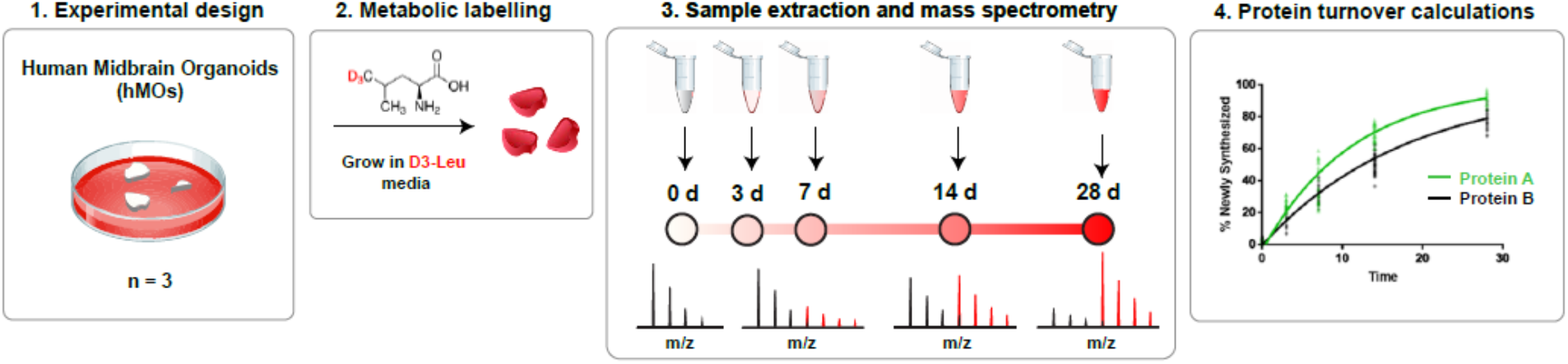
Experimental overview. Human midbrain organoids derived from healthy iPSC lines were metabolically labelled with D3-Leu SILAC media over a time course experiment. The organoids were processed at 5 different time points to extract proteins and digest into peptides for MS analysis at each time-point. Protein identifications and the rate of heavy isotope incorporation was determined through MS. Specific turnover rates on a protein level were calculated using MaxQuant, Skyline, and Topograph (all open-source software).

**Figure 2.**
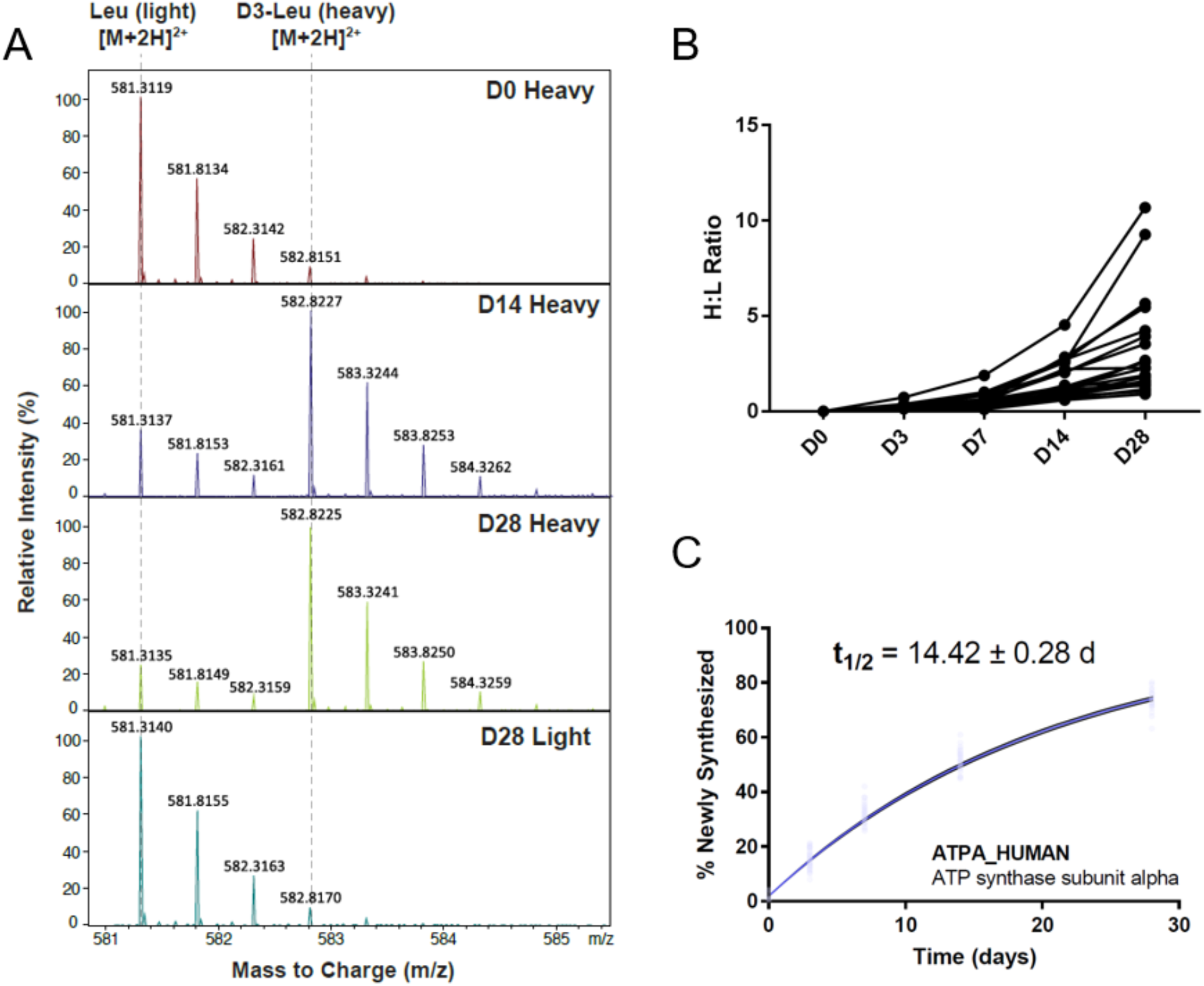
D3-Leu incorporates into hMOs and produces robust protein half-life calculations. **A**) A representative MS1 spectrum of a peptide at 3 different time points displaying successful incorporation of heavy D3-Leucine (H = heavy label peak, L = light label peak). B) Heavy to Light (H:L) ratios were calculated on all identified hMO proteins using MaxQuant. A progressive increase of H:L ratio can be observed over the time course. C) Example of Topograph half-life output for ATP synthase subunit alpha. Blue dots represent the 23 peptides across all the time points that contributed to the half-life calculation. Black lines indicate the 95% confidence interval.

Overall, a total of 3280 proteins derived from 20842 peptides were identified from our MS data. After removing peptides that did not meet the acceptance criteria (2.6.4), 844 proteins remained for analysis. All hMO protein half-lives were summarized and grouped according to KEGG annotation or cellular localization to investigate trends in compartment- or function-specific turnover rates (Figure 3). hMO half-lives ranged from 2 to 15 days, with an average half-life of 9.16 days. Most annotated protein groups did not deviate significantly from the population average, except for a few notable exceptions: first, mitochondrially localized proteins displayed significantly longer half-lives compared to all measured hMO proteins. Second, histones, proteasomal subunits, and ribosomes were also significantly longer lived. Long histone lifespans could be necessary for maintaining chromatin structure [35]. These findings also align with previous studies showing histones exceptional stability and persistence in mammalian models [22,35]. Protein groups that demonstrated shorter lifetimes (compared to the rest of hMO proteins) consisted of those involved in endocytosis and RNA transport.

**Figure 3:**
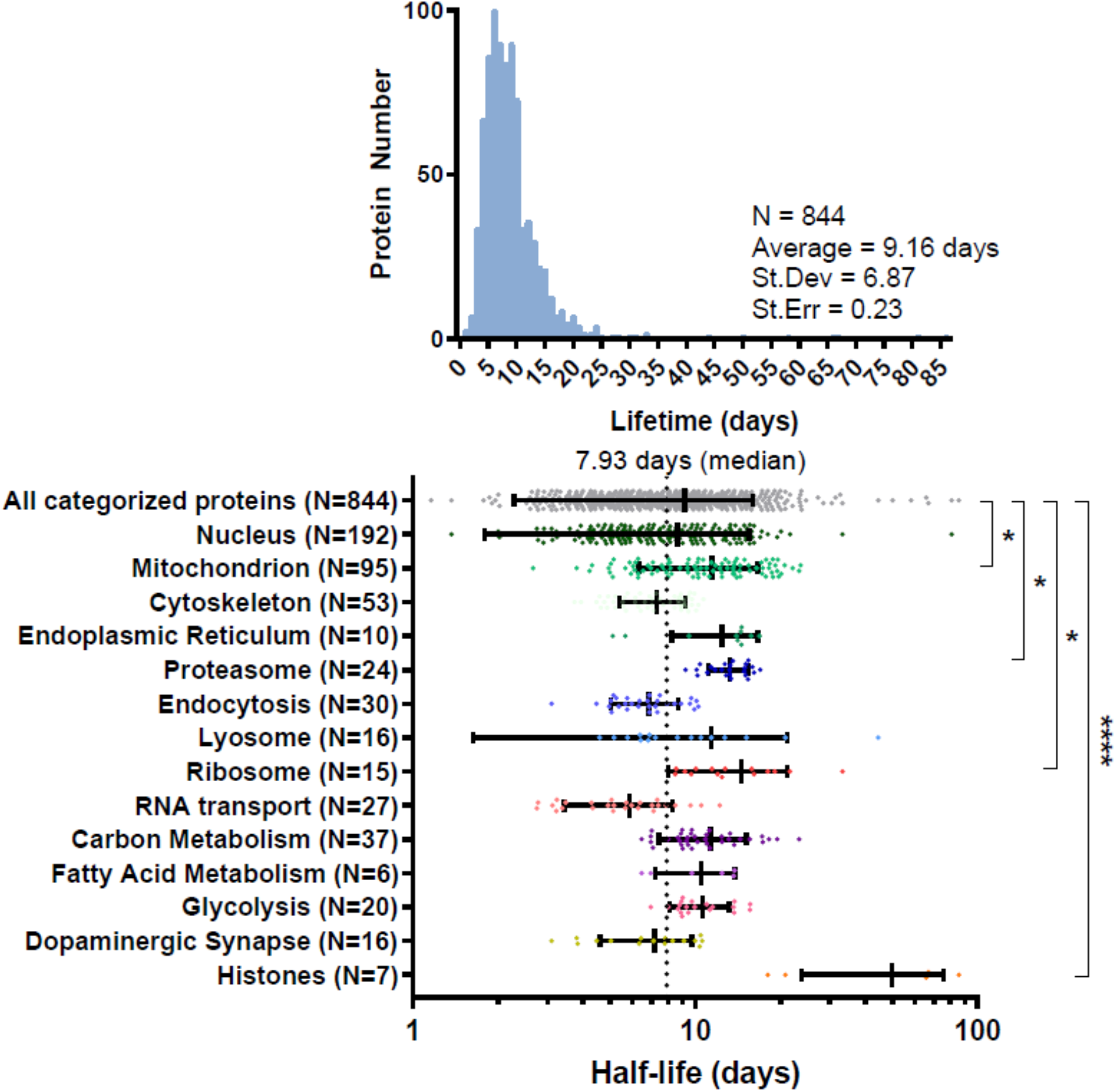
Lifetime of organoid proteins. Lifetimes of proteins organized into groups accordingly to their location and biological pathway. Each data point corresponds to a single protein lifetime, derived from 3 biological replicates and at least 2 distinct peptides. The black lines indicate the mean and the standard deviation (SD) for each group. Analysis of variance (ANOVA) followed by the Dunnett’s test summarizes the significance of protein groups compared to the average half-life of all the proteins identified (* ≤ 0.05, **** ≤ 0.0001).

## 3.2 Comparison of turnover rates with prior studies

This is the first time that protein half-lives have been characterized in brain organoids. As such, we sought to compare our data with previously published turnover measurements in mice [22]. Overall, proteins were turned over faster in organoids than in mice (organoid_t1/2_ = 9.16 days vs. mouse_t1/2_ = 10.7 days, P < 0.0001). When compared to different organs and brain regions in mice, hMOs were most similar to mouse hindbrains, albeit modestly (Figure 4A). To our knowledge, there are no datasets measuring protein turnover in mouse midbrains, so a direct comparison is not yet feasible. Still, the relative turnover of protein groups seemed to be conserved across organisms, as mitochondrial proteins were longer lived than all other proteins for both mouse and hMOs. Next, we compared our data to two separate studies measuring turnover in primary cultured neurons (Figure 4B) [36,37]. In both cases, the average lifetime of proteins in 2D-cultured neurons were faster than that of organoids. Taken together, our results position hMOs as a model of mammalian proteostasis that lies between an *in vitro* 2D cell culture and an *in vivo* system. Future studies will be critical in profiling protein turnover within different organoid models to confirm these findings across tissues. Still, it will also be essential to clarify and resolve some inherent limitations in the current methodology for SILAC-based measurements in organoids.

**Figure 4:**
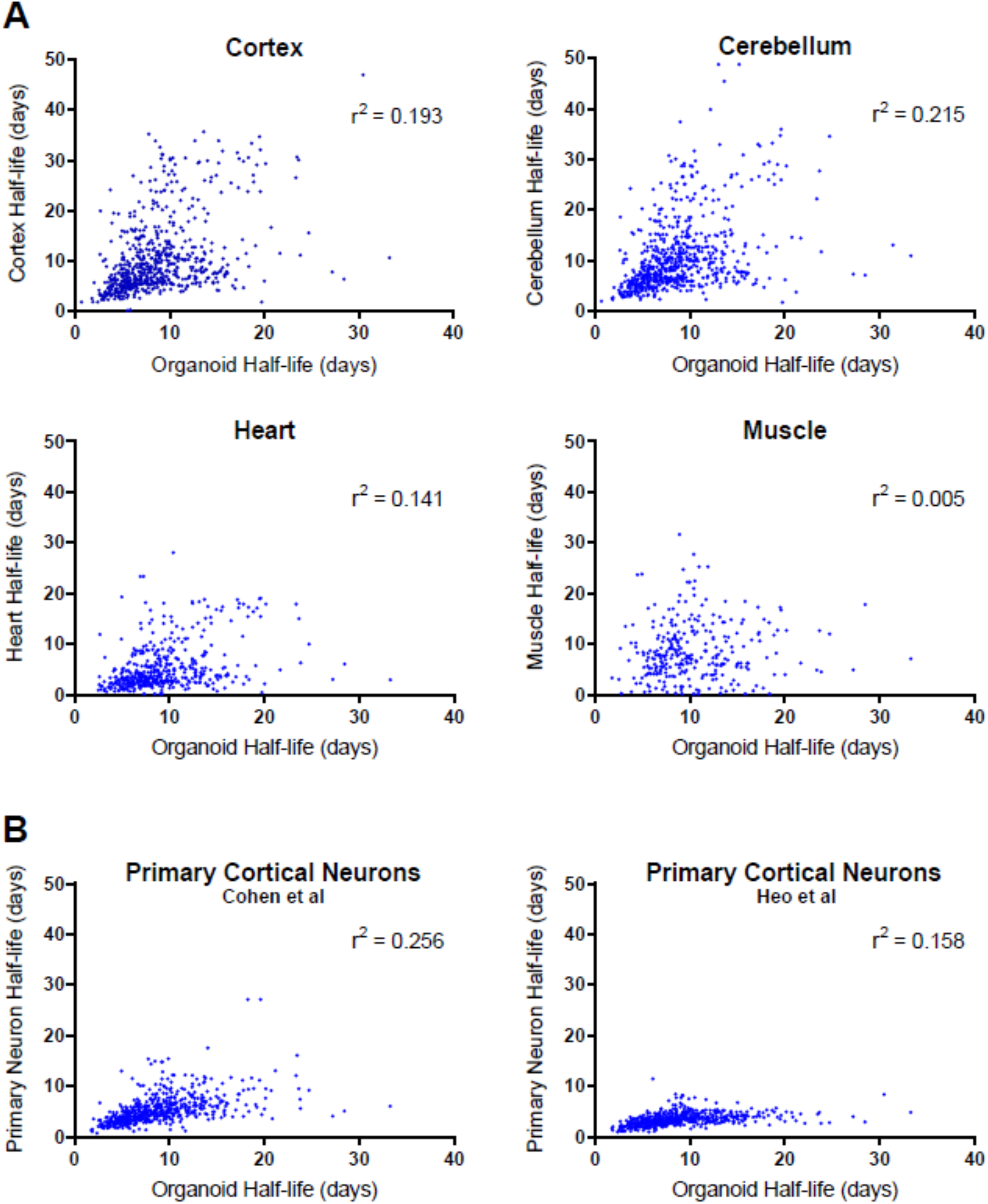
Correlation between global hMO half-lives with mice and primary cultured neurons. A) Scatterplot comparing lifetimes of proteins *in vivo* of four different organs with our organoid data, with Pearson’s correlation coefficients denoted by r^2^. B) Scatterplot comparing lifetimes of proteins with *in vitro* data from rat cortical neurons [36,37]. Correlation was determined through matching the corresponding datasets with our organoid proteins by gene name.

## 3.3 Limitations

Organoids have immense potential for the modelling and understanding of various diseases, but still suffer from important limitations. Notably, organoids lack an efficient circulatory system leading to issues with oxygen and nutrient exchange [38]. Likewise, due to the inherent 3D organization of the tissue culture, cell necrosis has been observed at the core of many reported organoid models, which may contribute to a background pool of unlabeled proteins in our system [2]. Recent work has highlighted the utility of microfluidic devices in organoid systems to improve nutrient access and reproducibility in growth, which may also facilitate future studies in hMOs [39]. It is also important to note that turnover measurements in their current form necessitate a few key assumptions: (1) The biological system must be in a steady state; this can be verified by quantifying the total level of protein at the beginning and the end of the time course, as we did here (2) All fragments of a protein are turned over at the same rate (i.e. one turnover measurement is calculated for each protein, even if a proteolytic fragment or domain of a protein may be turned over more rapidly). Finally, it is also important to note that brain organoids are small and thus axons do not grow more than a few millimeters, whereas axons in mammals can be several centimeters, and even meters. This could create significant differences in the turnover rate of proteins located in the soma compared to the synaptic terminals. Thus, the apparent differences between our lifetime calculations and previous studies could also reflect limitations with hMOs and our current experimental set-up.

## 4. Conclusion

We have provided a robust framework for extracting proteins and measuring global protein turnover in human midbrain organoids. We have also developed a simple analytical and statistical workflow that can be implemented by scientists of all skill levels using open-source, freely available software. Using this methodology, we have shown that human midbrain organoids have a global protein turnover that is faster than mice, but slower than 2D neuronal cultures. Future work using our approach will be able to highlight crucial differences in protein turnover between control and disease models of brain organoids. Overall, our work facilitates the study of proteostasis in organoid models of human disease and will provide a framework to measure protein turnover in organoids of all cell types.

## Acknowledgments

We thank the McGill Pharmacology SPR/MS facility (M. Hancock) and the CFI for support, as well as the Proteomics platform at the Research Institute of the McGill University Health Centre (Lorne Taylor, Amy Wong, Jennifer Nedow). This work was supported by an *Innovation Ideas* grant from the *Healthy Brains, Healthy Lives* program at McGill. J.D and A.D were supported by a *Canada Graduate Scholarship* from the Canadian Institutes for Health Research (CIHR), and J.FT. holds a Canada Research Chair (Tier 2) in Structural Pharmacology (CIHR). E.A.F. is supported by a Foundation grant from the CIHR (FDN-154301) and a Canada Research Chair (Tier 1) in Parkinson’s Disease.

